# An open protocol for modeling T Cell Clonotype repertoires using TCRβ CDR3 sequences

**DOI:** 10.1101/2022.03.30.486449

**Authors:** Burcu Gurun, Wesley Horton, Dhaarini Murugan, Biqing Zhu, Patrick Leyshock, Sushil Kumar, Katelyn T. Byrne, Robert H. Vonderheide, Adam A. Margolin, Motomi Mori, Paul T. Spellman, Lisa M. Coussens, Terence P. Speed

## Abstract

T cell receptor (TCR) repertoires can be profiled using next generation sequencing (NGS) to monitor dynamical changes in response to disease and other perturbations. Several strategies for profiling TCRs have been recently developed with different benefits and drawbacks. Genomic DNA-based bulk sequencing, however, remains the most cost-effective method to profile TCRs. The major disadvantage of this method is the need for multiplex target amplification with a large set of primer pairs with potentially very different amplification efficiencies. One approach addressing this problem is by iteratively adjusting the concentrations of the primers based on their efficiencies, and then computationally correcting any remaining bias. Yet there are no standard, publicly available protocols to process and analyze raw sequencing data generated by this method. Here, we utilize an equimolar primer mixture and propose a single statistical normalization step that efficiently corrects for amplification bias post sequencing. Using samples analyzed by both approaches, we show that the concordance between bulk clonality metrics obtained from using the commercial kits and that developed herein is high. Therefore, we suggest the method presented here as an inexpensive and non-commercial alternative for measuring and monitoring adaptive dynamics in TCR clonotype repertoire.

## INTRODUCTION

The receptors on the surface of T cells bind to an enormous array of antigens that play a pivotal role in shaping immune response during health and disease. The T cell receptor (TCR) is a heterodimer composed of one alpha and one beta chain which are encoded by the *TCRα* and *TCRβ* genes, respectively. To recognize an extremely large antigen space, the TCR genomic loci undergo somatic recombination of variable (V), diversity (D), and joining (J) gene segments, and generate a diverse repertoire of TCRs. The complementarity determining region 3 (CDR3) region present at the D segment of the recombined *TCRβ* gene is highly diverse in TCR beta chains. Therefore, surveying the recombined *TCRβ* gene or transcript as a proxy for overall TCR repertoire diversity has emerged as a rational approach to study TCR repertoire dynamics.

Over the last ten years it became possible to obtain comprehensive profiles of TCR through an array of next generation sequencing (NGS) based approaches (1-8). These vary based on the experiment type (bulk or single-cell), sample type (RNA or DNA), library preparation (multiplex PCR, bead-based enrichment, 5’RACE) and sequencing platform, each choice presenting a different, and fascinatingly interlocked trade-off. Single cell approaches compared to bulk can be very accurate and unbiased for high frequency clones, but have lower resolution for low frequency clones (9,10). They are also substantially more expensive and require intact cells. RNA-based approaches are affected by the variability in TCR RNA expression levels, but may better reflect diversity when the sample size is limited (8). Genomic DNA (gDNA)-based approaches require either multiplex PCR or target enrichment during library preparation which introduces biases (11). A recent comparison of these two approaches confirmed these issues for both RNA- and DNA-based methods but also found the methodological variability to be smaller than the biological variability (12). New innovations are being introduced rapidly for all of these approaches but currently there is no established gold standard.

Multiplexed PCR-based bulk sequencing approaches using gDNA, however, have become the standard approach for translational and even clinical applications due to reasonable sample requirements and moderate costs (13). This is reflected in the fact that all currently available commercial TCR sequencing products offer this as their major DNA option (Adaptive Biotechnologies (14), BGI (15), iRepertoire). Multiplex PCR refers to the usage of multiple forward primers specific for the V segments and multiple reverse primers specific for the J segments in combination during the initial amplification for target enrichment. Since each primer pair will have a different efficiency multiplexing will distort the relative abundances of the VDJ segment combinations (16). Correcting for this amplification bias is a key challenge for accurate quantification. One approach (14) is using spiked-in oligomers (synthetic templates) for each primer pair to measure primer efficiencies and to carefully control the design of the primers and their concentrations accordingly. Assuming that there is no interaction between the efficiency of primer pairs, amplification bias is reduced by iteratively calibrating the primer concentrations to find the optimal primer mix, and then removing any remaining amplification bias computationally using spiked-in oligomer counts.

The proprietary setup described in (14) is currently available for human and murine samples exclusively through commercial kits. These are difficult and labor-intensive to adapt if requiring different settings, e.g. a different mammal or application. Furthermore, synthetic templates when added to all of the samples with the kits can substantially increase preparation and sequencing costs, as well as decrease coverage for clonotypes.

Here we demonstrate, that using a negative binomial mean normalization strategy, it is possible to normalize amplification bias within a reasonable error margin without balancing primer concentrations using synthetic templates. Furthermore, we report that amplification bias parameters are highly preserved between datasets. This observation indicates that one can use synthetic template normalization controls for a small subset of samples, which can then be used to calculate mean scaling factors per batch/experiment, thus substantially reducing the cost and coverage of TCR repertoire analysis.

## MATERIAL AND METHODS

### Multiplex primers and design of synthetic TCR templates

Multiplex PCR primers previously described by Faham *et al*. (US patent 8,628,972 B2), for the amplification of murine *TCRβ* genomic loci were utilized **(Supplementary Table 1: primer sequences)**. The 20 Vβ segment specific primers amplify all the 21 functional Vβ segments, and the 13 Jβ specific primers amplify all the 13 functional Jβ segments. As previously described by Carlson et al., we designed 260 (20V x 13J) synthetic TCR templates (ST) to minimize amplification bias due to multiplexing with 20 Vβ forward and 13 Jβ reverse primers (14). Briefly, ST are 200 bp long double stranded DNA segments that contain partial V segment and J segment sequences encompassing a set of internal barcodes for post-sequencing identification. The internal barcode region contains a 16 bp barcode specific for each VJ combination. This specific barcode is further flanked by a 9 bp barcode that is common for all ST. An equimolar mixture of the 260 ST is added to the genomic DNA samples during PCR as internal controls.

### Amplification and deep sequencing of TCRβ genomic locus

Genomic DNA from freshly resected mouse peripheral blood, peritoneal orthotopic mesotheliomas (17), and spleen was isolated using the Qiagen DNeasy Blood and tissue kit. Utilizing the method described in Robins et al (18), we performed a 2-stage PCR using genomic DNA for TCRβ deep-sequencing library preparation. The 1^st^ stage involved amplifying the gDNA and the ST using a multiplex PCR with 20 Vβ forward and 13 Jβ reverse primers using the Qiagen multiplex PCR kit. The multiplex PCR primers contain a common 5’ overhang, allowing amplification by a single primer pair in the 2^nd^ stage PCR. Using 2.0% of the purified PCR product from stage 1 as template, a 2^nd^ stage PCR was performed with universal and indexed Illumina adaptors. Of note, the indexed adaptors contained an 8-base index sequence, providing each sample with a unique sample barcode. Equal volumes of all samples were pooled. Each pool concentration, typically containing PCR mixtures from 70 samples, was measured with a 2200 TapeStation (Agilent), and the concentration determined by real time PCR using a StepOne Real Time Workstation (ABI/Thermo) with a commercial library quantification kit (Kapa Biosystems). Paired-end sequencing was performed with a 2 × 150 protocol using a Midoutput 300 sequencing kit on a NextSeq 500 (Illumina). Target clustering was ∼ 160 million clusters per run. Following the run, base call files were converted to fastq format and demultiplexed by a separate barcode read using the most current version of Bcl2Fastq software (Illumina).

### TCR data analysis Pipeline

Fastq files were assessed for initial read quality using the FASTQC public tool (19), including the per-base quality scores. Quality paired-end sequences were combined using the PEAR (Paired-End reAd mergeR) algorithm (20). Merged sequences were then separated into ST and non-ST sequences. ST sequences were identified by searching for the common flanking 9-bp internal barcodes allowing a one-nucleotide mismatch or indel. Sequences flagged as ST via this search were removed from downstream clonotype analyses. The individual ST sequences were distinguished and quantified by searching for the specific 16-bp barcode sequences unique to each ST, again allowing a one-nucleotide mismatch or indel **(Supplementary Figure 1)**. Clonotypes were identified from purified (ST-removed) sequences utilizing the MiXCR pipeline (21), which is a two-step alignment and assembly process. First, reads were aligned to reference V, D, and J sequences, using the align module. Next, the assemble module grouped alignments into distinct clonotypes using a hierarchical clustering method based on sequence similarity and relative abundance. Finally, the export module exported alignments as well as assembled clones in tabular format. Raw clonotype counts were normalized using the NB mean normalization strategy described below. Normalized clonotype counts were exported in tabular format for use in downstream analysis. A number of TCR repertoire metrics, including clonality, maximum clonal frequency, and the Shannon diversity index were calculated. Quality control data was recorded in an overall summary table, for reference **(Supplementary Material 1)**.

### Data Sets

#### ST-only data sets

Twenty samples of an equimolar mixture of ST were sequenced. Two sets of ten samples of equimolar mixtures of ST at different concentrations were also sequenced at a later date. No genomic DNA was present in these samples.

#### Transgenic TCR data sets

A dilution series was created from spleen genomic DNA of P14 and OT-1 TCR transgenic mice (C57BL/6-Tg(TcraTcrb)1100Mjb/J). Three replicates of 300, 600, 900, and 1200 ng of DNA from P14, OT1, and a 50:50 mixture of P14 and OT1 DNA were sequenced along with an equimolar mixture of ST for a total of 12 samples. Appropriate amplification of the transgenic clones was assessed. (**Supplemental Figure 2**).

#### ST dilution series data set

A dilution series was created from ST at six levels as below:

* Stock: 3.2 ng/ul (equimolar mixture of all 260 spikes)

Dil1= 0.1 of stock

Dil2= 0.1 of Dil1

Dil3= 0.01 of Dil1

Dil4= 0.01 of Dil3

Dil5= 0.001 of Dil3

Dil6= 0.001 of Dil5

Three replicates of 600 ng of blood DNA of murine were added to all levels of dilutions and for no ST samples for a total of 21 samples and three replicates of 600 ng of tumor DNA of murine were added to all levels of dilutions and for no ST samples for a total of 21 samples.

#### WT spleen data set

Wild-type mouse spleen genomic DNA were sequenced for a total of five replicates.

#### Byrne et al. data set (22)

gDNA from 17 murine pancreatic ductal adenocarcinoma specimens were previously sequenced by a commercially available TCR beta platform. Aliquots of these gDNA samples were obtained and sequenced along with an equimolar mixture of ST using the protocol described above.

#### Mesothelioma data set

Peritoneal mesothelioma tumors derived from the 40L cell line (17)derived genomic DNA samples derived from 12 syngeneic mice were sequenced along with equimolar mixture of ST.

### Batch mean scaling factors

This method creates a *scaling factor* for each ST (and therefore for each primer pair) based on that ST’s counts among all samples, relative to all ST counts within a batch. Given a matrix (*C*_*ij*_) of ST counts from a batch, where *i=1,…,260 labels* ST and *j=1,…,n labels* samples in the batch, we denote the batch mean of the counts for ST *i* by *C*_*i•*_ = *n*^−1^ ∑_*j*_ *C*_*ij*_ and the batch mean of all ST means by *C*_••_ = (260)^−1^ ∑_*i*_ *C*_*i*•_ = (260*n*)^-1^ ∑_*i,j*_ *C*_*ij*_. The scaling factor (SF) for ST *i* in that batch is *SF*_*i*_ = *C*_*i*•_⁄*C*_••_.

### Negative Binomial means

The above idea of a scale factor is distribution free, but for its use in normalizing counts, would require a full set of ST in every sample. We explored the use of a ST-specific negative binomial model to dispense with the use of synthetic templates. Consider the set *(C*_*1*_, *…*., *C*_*n*_) of counts for single, fixed ST across a batch of *n* replicate ST-only samples. A plausible model for these counts is the negative binomial (NB) distribution. We write *C* ∼ *NB*(*m, d*) for this distribution, where *m>0* is the *mean* parameter and *d≥0* is the (over-) *dispersion* parameter, and refer to (23) for an explicit formula for the *NB* probability mass function. For present purposes, *S* has expected value ***E***(*C*) = *m* and variance ***var***(*C*) = *m*+ *dm*^2^. When *d=0*, the negative binomial reduces to the Poisson distribution, for which ***E***(*C*) = ***var***(*C*), and thus the use of the term over-dispersion here is relative to the Poisson. Using the methods of generalized linear models (23), we can obtain the maximum likelihood estimates (MLE) 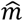 and 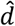 of *m* and *d* from a replicate set of ST such as *(C*_*1*_, *…*., *C*_*n*_). In the notation of the previous paragraph, the MLE 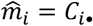, that is, the MLE of the *i*th mean parameter *m*_*i*_ of an NB fitted to *(C*_*ij*_) is the arithmetic mean of the *i*th set of observed counts (assumed independent and identically distributed across *j=1,…,n* with common mean *m*_*i*_ and common dispersion parameter. Where no confusion will result in what follows, we will not distinguish the parameters *(m*_*i*_) from their (maximum likelihood) estimates 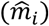. These ST or primer-pair-specific means estimated from ST-only data can be used as scaling factors for normalization, even when there are no ST present in samples. Consistent with the notation in the previous paragraph, we write *m*_•_ = (260)^-1^ ∑_*i*_ *m*_*i*_ for the average of the 260 (estimated) mean parameters. The 260 NB mean scaling factors are (*m*_*i*_⁄*m*_•_).

### Normalization

To normalize a set of clonotype counts from a single sample, we first calculated the primer-pair totals *(C*_*i*_), where *C*_*i*_ denotes the total count of all clonotypes amplified with primer-pair *i*, where *i=1,…,260*. We then normalize the 260 counts *(C*_*i*_) using the batch scaling factors *(SF*_*i*_) by dividing by the corresponding scaling factor: 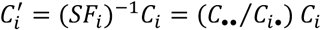. Similarly, we normalize the *(C*_*i*_) using the (estimated) NB mean scaling factors (*m*_*i*_/*m*_•_) by dividing: 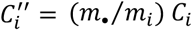. After these primer-pair totals were normalized, the counts for distinct clonotypes sharing the same primer-pair were normalized: if one such clonotype accounts for a proportion *p* of the total count *C* corresponding to its primer pair, then it will be assigned a normalized value equal to the same proportion *p* of the normalized primer-pair total *C’* or *C”*.

The same normalization could be used for ST counts if available. That is, divide the observed count of the fragments arising from primer pair *i* by the *SF*_*i*_ or *m*_*i*_⁄*m*_•_ for primer pair *i* for both ST and clonotype counts alike. Since the mean count of ST *i* will be proportional to *m*_*i*_, normalization should preserve the total count of ST, exactly for batch mean normalization, on average for *NB* normalization. As long as the observed clonotype counts exhibit the same relative over- and under-representation after amplification as that exhibited by the ST, any bias will be reduced by this normalization.

## RESULTS

### Amplification bias due to multiplex PCR is reproducible

To amplify all possible TCR somatic recombination products, we performed a multiplex PCR with 20 different V-specific forward primers and 13 different J-specific reverse primers. Since the differences in efficiency of primer pairs can produce significant amplification bias in TCR clonotypes, we spiked-in 260 ST in equimolar concentration as internal controls to the multiplex PCR reaction in order to measure and control bias (**Figure 1A**). We hypothesized that the estimated mean counts of the 260 ST would scale proportionally across samples and experiments. To test this, we measured the distribution and variation of ST counts across 20 ST-only samples in the absence of genomic DNA **(Figure 1B)**. Within each sample, ST counts were converted to relative frequencies to reduce the effect of random sample-to-sample variation on the comparisons. For a given ST, deviation of the median relative frequency across all samples from 1/260 is a measure of the PCR amplification bias for the corresponding primer pair, while the spread of relative frequencies is an indication of the random sample-to-sample variation. We observed that the ST-to-ST variation was much larger than the sample-to-sample variation within an individual ST (∼20-fold difference in respective median IQRs). These differences indicate that the observed differences in ST counts are primarily caused by amplification bias of different primer pairs. The ln (1/260) line indicates the (log) expected ST relative frequency in the absence of amplification bias. To demonstrate that the above observations apply in the presence of TCR clonotypes, we obtained ST counts from samples with genomic DNA extracted from P14 and OT-1 TCR transgenic mice where CD8 T cells primarily recognize OVA_257-264_ when presented by the MHC I molecule. A similar relationship was found between ST relative frequencies in the presence of TCR clonotypes across samples, though the sample-to-sample variation is noticeably larger **(Supplementary Figure 3)**.

**Figure 1.**
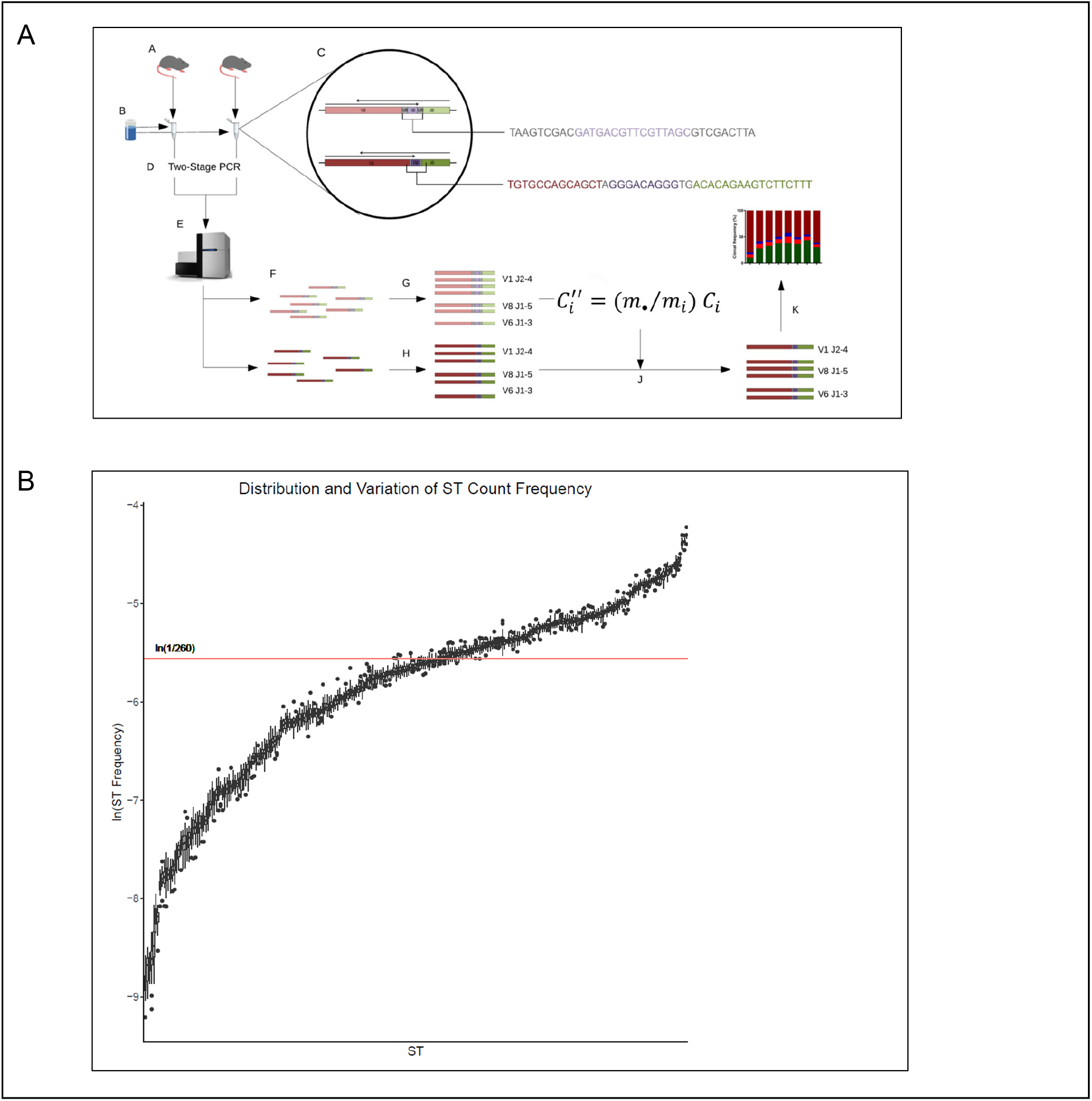
Pipeline overview and ST proportion distributions. (A) Overview of the TCRβ seq analysis pipeline: **A**. gDNA extracted from freshly resected murine peripheral blood, peritoneal orthotopic mesotheliomas, and spleen tissues. **B**. Equimolar mixture of synthetic TCR templates (ST) were then added, and **C**. followed by addition of forward and reverse sequences of ST (top) and TCRβ (bottom). For ST, universal 9-bp barcode (grey), unique 16-bp barcode (purple). For TCRβ, V-region (red), VD/DJ-junctions (grey), D-region (purple), J-region (green). **D**. Samples amplified with multiplex PCR followed by second-stage barcoding PCR. **E**. Samples were then pooled for sequencing. **F**. ST and TCRβ were then separated using universal barcodes, and **G**. ST was quantified using unique barcodes. **H**. TCRβ clonotypes were quantified with the MiXCR tool suite. **I**. Negative binomial normalization was used to remove amplification bias. **J**. Scaling factors were applied to counts, and **K**. then used to normalize counts for diversity analyses. (B) Plot showing stability for relative frequencies of ST counts within individual samples, and amplification bias based on the reproducibility of the multiplex PCR. Median IQR of ST-to-ST variation was ∼20-fold greater than the median IQR of experiment-to-experiment relative frequencies of ST counts. Target values aim to be at the ln (1/260) line in the absence of amplification bias. Data derived from twenty ST-only samples described in the *ST-only data sets*.

### A negative binomial model fits the data

The fit of count data on 260 ST from each of 20 ST-only samples to the negative binomial (NB) with ST-specific means and a common dispersion parameter *d* can be informally assessed by examining an empirical variance (*v*) vs mean (*m*) plot with the line *v = m + dm*^*2*^ displayed, all with log-log scales.

At the high end, we expect an approximate linear relationship between log mean and log variance for the ST counts, with a slope of about *2*, since *log v ≈ log d + 2log m* for large m. In **Figure 2**, we took *d=0*.*125*, as this was the median value of the dispersion estimates found by fitting separate NB distributions to the 260 sets of 20 ST counts.

**Figure 2.**
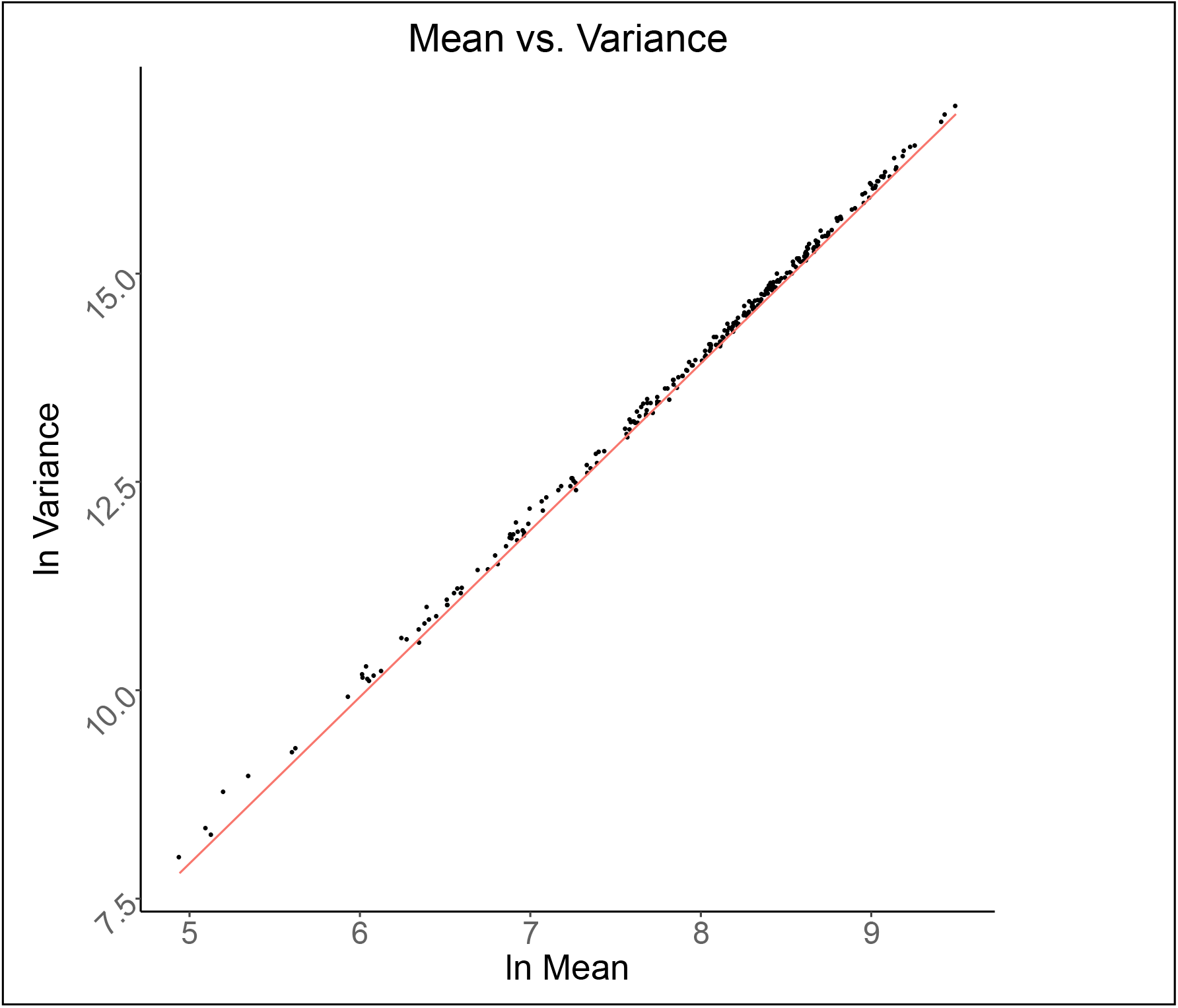
A negative binomial model fits the data. Observed mean-variance relationships fit as compared to the negative binomial (NB) model. Dots are observed values derived from 260 ST count distributions with a separately fitted dispersion parameter. The red line is the relationship predicted by the NB model with fixed d (0.125). Data derived from twenty ST-only samples described in the *ST-only data sets*.

The fact that the ST counts fit NB distributions with approximately similar overdispersion parameters reassures us the variation we are seeing is in some sense natural and that the system is in control(24).

### Scaling factors are relatively stable across experiments

We fit the NB model to three different sets of ST-only observations at different equimolar concentrations, coming from two different batches: (1) a set of 20 samples from one batch, (2) two sets of 10 samples from another batch. To demonstrate that scaling factors were relatively stable across ST-only samples, we compared the sets of 260 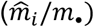 values based on 20 ST-only samples, to those based on the sets of 10 ST-only samples. We observed the 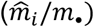 values to be highly correlated on a *log-log* scale (Pearson *r* = 0.84 and 0.97, *p*-value < 2×10^−16^ for both comparisons) (**Figure 3A-B)**, indicating that scaling factors are relatively stable across experiments.

**Figure 3.**
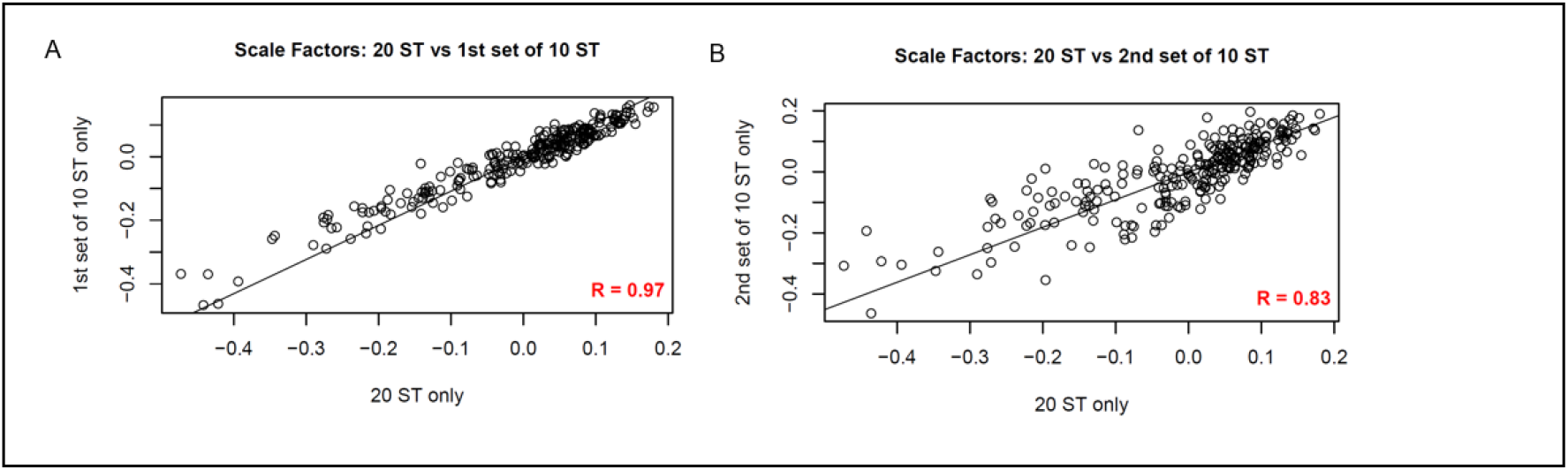
Stability of scaling factors. (A, B) Scatter plots (*log-log* scales) comparing the sets of 260 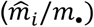 values based on 20 ST-only samples to those based on the two sets of 10 ST-only samples (described in the Data Sets section). The 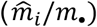 values are observed to be highly correlated across different batches and samples. (Pearson r = 0.97 and 0.84, p-value < 2×10^−16^ for both comparisons).

We then computed estimates of the (*m*_*i*_) by pooling data across these sets, adjusting for the concentration differences, calling the results the combined estimates. Pooling also deals with errors introduced to the ST-only counts by processing samples in different batches along with different samples. The combined estimates of the (*m*_*i*_) are therefore used for normalization below (**Supplementary Appendix 1**).

### Normalization considerably reduces amplification bias

We normalized the 20 ST-only measurements with the combined estimates of the (*m*_*i*_). The spread of the ST counts for 20 samples was considerably reduced after normalization **(Figure 4A)**. A similar comparison of ST counts for 20 samples, in the presence of genomic DNA derived from mesothelioma tumors, revealed that the reduction of ST counts was present, although less pronounced **(Supplementary Figure 4)**. To validate the normalization procedure further, we assessed the observed ratio of monoclonal counts before and after normalization utilizing the 50:50 mixture of P14 and OT-1 TCR transgenic monoclonal DNA. After normalization, the differences between the transgenic TCR counts were substantially reduced for all samples, as represented by the smaller deviations of the proportion of the dominant clonotype from ½ following normalization (**Figure 4 B)**.

**Figure 4.**
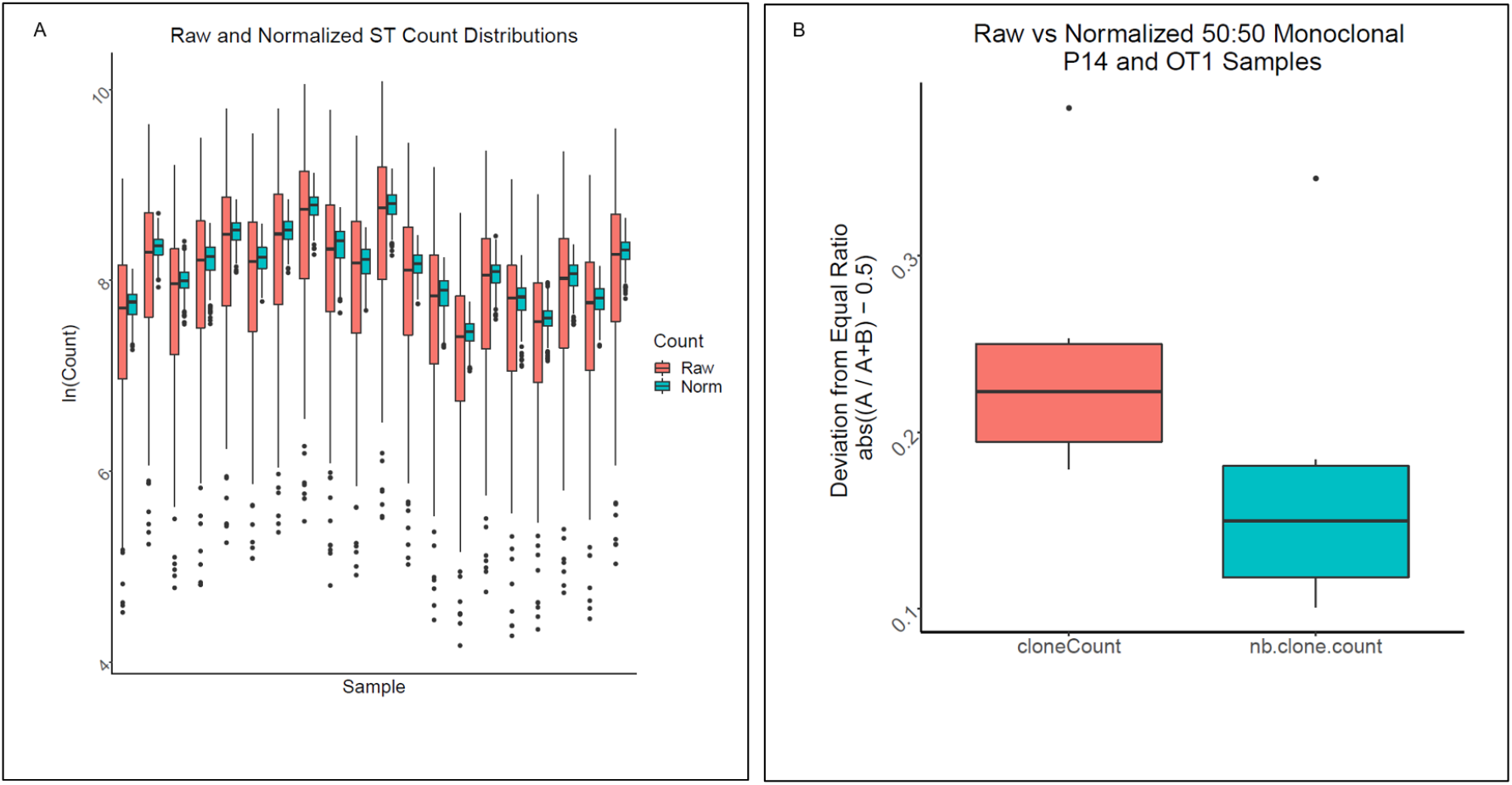
Normalization reduces amplification bias. (A) NB normalization reduces amplification bias. Variation in ST counts within samples is apparent before normalization (red), however, the ST-to-ST differences within each sample are reduced to less than two-fold after normalization (cyan). Data derived from twenty ST-only samples described in the *ST-only data sets*. (B) Amplification bias of spleen genomic DNA of P14 and OT-1 TCR transgenic mice. To further evaluate normalization parameters, a 50:50 mixture of P14 and OT1 TCR transgenic monoclonal DNA was utilized to examine differences between transgenic TCR counts (red) that were reduced for all samples following normalization (cyan), as observed by the smaller deviations from ½ of the proportions of the dominant clonotype. Red color indicates values from clone counts before normalization, green color indicates values from normalized clone counts. Data derived from 12 50:50 mixture of P14 and OT-1 transgenic mice samples described in the *Transgenic TCR data sets*.

### Amplification bias reduction benefits from the dependence of primer pairs

A question from data presented in **Figures 4A and 4B** that arose had to do with determining how great a reduction in the spread of the 260 ST counts would be possible, given the variation, even in the absence of amplification bias. A theoretical analysis is presented in **Supplementary Appendix 2** under the assumption that counts from equimolar concentrations of the 260 ST are *independent* NB distributions with the same mean and dispersion parameters gives a lower bound. Seeing that counts from the 20 ST-only samples were well approximated by NB distributions with the same dispersion parameters (albeit with quite different means), our conclusion was that the different ST counts were likely not independent. Further, this result supports the normalization scheme, as the dependence aids reduction of the amplification bias below the level that would be expected under independence. Since each primer pair shares one primer with 32 other primer pairs, it is not surprising that the different ST counts are not independent; indeed, patterns of dependence in counts using chi-squared statistics are observed (**Figure 5**). The interactions revealed as patches of red and blue colors demonstrate that groups of V primers exhibit positive or negative dependence together with groups of J primers, that is, they interact to become over or under-represented in groups.

**Figure 5.**
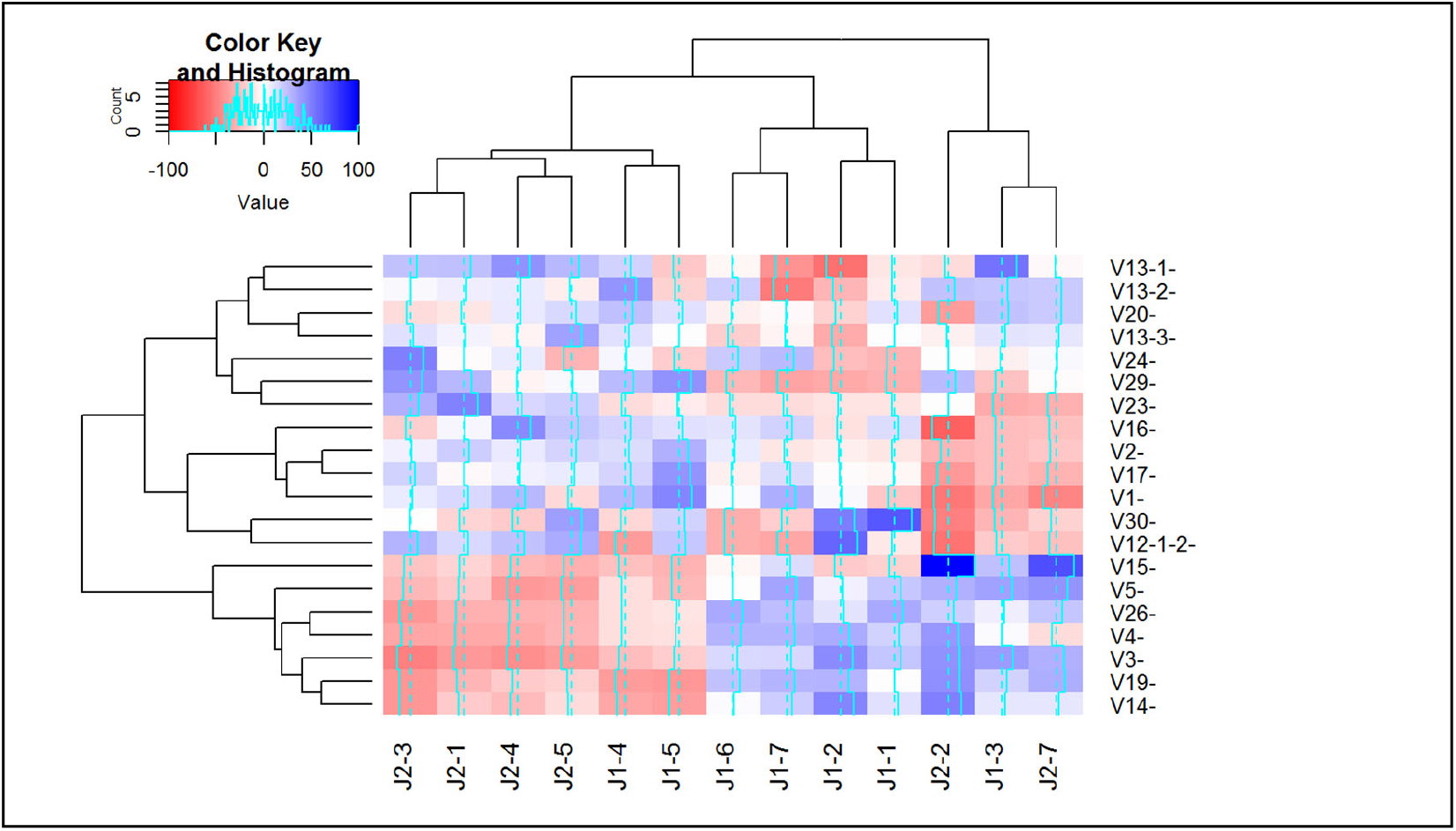
Cluster analysis revealing dependence between forward and reverse primers. A heatmap of interaction terms between forward and reverse primers reveal widespread and reproducible deviations from expected efficiencies under independence. Interaction terms were calculated as signed Pearson residuals (O-E)/√E where O is the observed ST count and E is the expected ST count under independence, calculated as E = row total x column total / grand total. Blue and red colors indicate positive and negative deviation from independence between forward and reverse primers respectively, and white indicates no deviation. Data derived from twenty ST-only samples described in the *ST-only data sets*.

In their experimental setup, Carlson et al., using an ANOVA based approach, concluded that primer pairs can be treated independently, and employed primer iteration experiments to find the optimal primer mix (14). Based on our normalization approach, however, the non-random, non-zero interaction terms revealed by chi-squared statistics substantially complicate preparation of an optimal primer mix through primer iteration experiments, and indicate a limit on the extent to which amplification bias can be addressed experimentally. We note that Carlson et al. also employed a second, computational normalization step, which we believe is primarily due to this limit.

### A negative binomial model supports downstream analyses of T cell repertoire dynamics

With the TCR clonotype counts normalized, we wanted to determine if results from these analyses were concordant with results from a commercially available platform (Adaptive Biotechnologies) described by Carlson et al (14). To achieve this, we utilized gDNA generated from pancreatic ductal adenocarcinoma specimens described in Byrne *et al(22)* containing 17 samples previously sequenced by Adaptive Biotechnologies’ platform. Aliquots of samples were sequenced and processed using the protocol described above. We utilized in-house software (see methods), tcR package, and VDJ tools to compute TCR repertoire metrics such as the diversity, clonality, and clonal distribution and refer to this pipeline as Open TCR Sequencing Protocol (OTSP). The 17 samples were sequenced in parallel and results were evaluated for concordance using Pearson correlation analyses for all combinations of: 1) Adaptive Biotech. platform sequences run by OTSP; 2) OTSP sequences run by OTSP; and 3) Adaptive Biotech. platform sequences run by Adaptive Biotech. **Figure 6A** and **Figure 6B** show concordance between results of OTSP sequences run by OTSP and Adaptive Biotech. sequences run by OTSP for Clonal index (*r*=0.8) and Shannon diversity index (*r*=0.8). The concordance for other combinations for Clonal Index and Shannon Diversity, as well as concordance between results of OTSP sequences run by OTSP and Adaptive Biotech. platform sequences run by Adaptive Biotech. for the frequency of hyperexpanded clones are shown in **Supplementary Figure 5 (A-E)**. Overall, results from the OTSP pipeline demonstrates a strong association with results from the Adaptive Biotech. TCR sequencing platform (p<0.001 for all comparisons).

**Figure 6.**
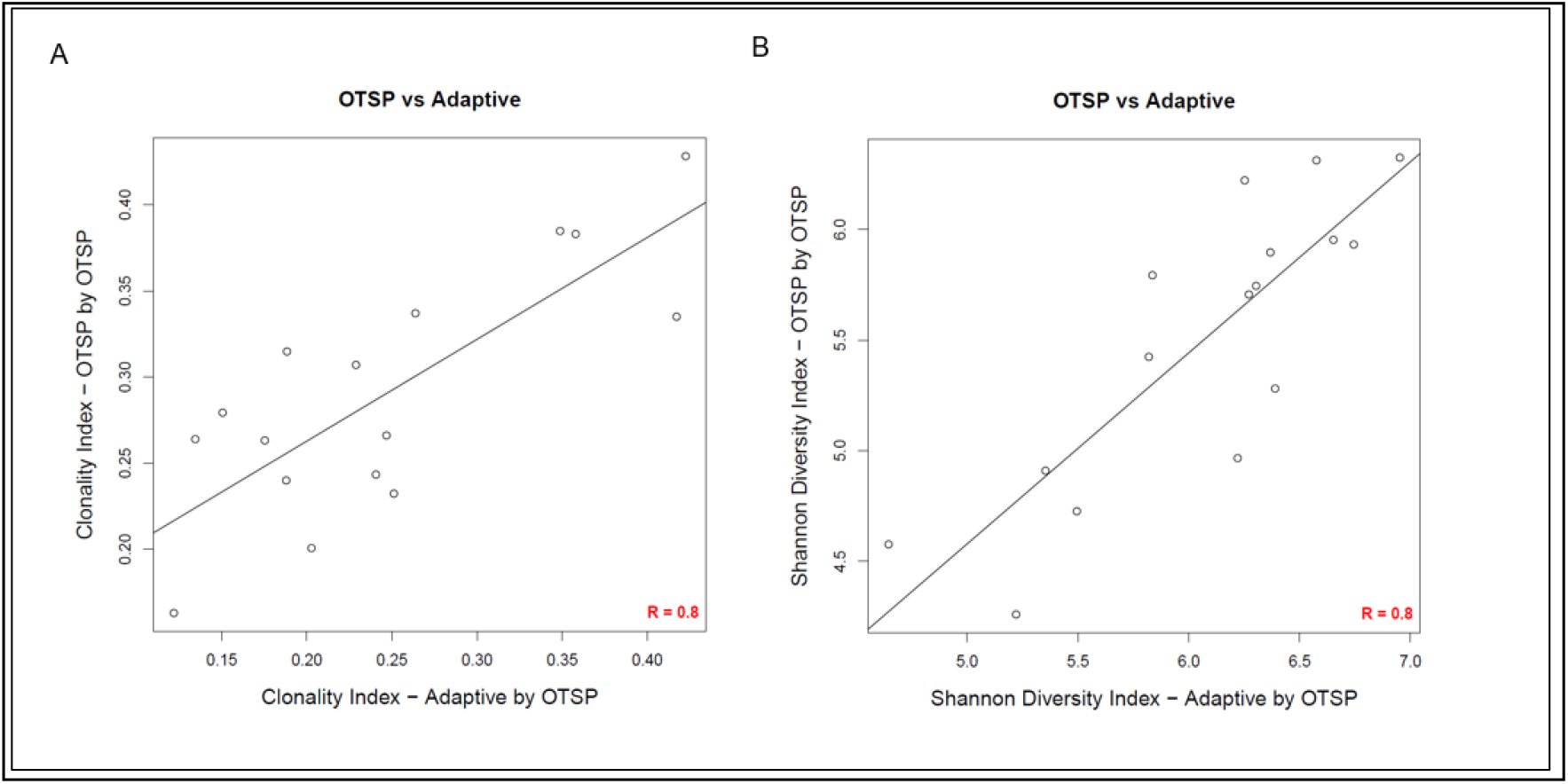
Concordance analysis of T cell repertoire metrics. Concordance analysis comparing commercial and in-house pipelines where samples were evaluated based on the results of OTSP sequences run by OTSP and the Adaptive Biotech platform sequences run by OTSP for Clonal Index (A) and Shannon Diversity Index (B). Pearson *r*=0.8 for both comparisons. (p<0.001). Data derived from samples described in the *Byrne et al. data set*.

The descriptions and formulas related to Clonal index, Shannon diversity index and frequency of hyperexpanded clones are included in (25), which deploys the NB mean normalization methodology described here in a biological context.

## DISCUSSION

Measuring and monitoring adaptive dynamics in patient TCR repertoires could have a significant impact on response and resistance monitoring for patients receiving various forms of immunotherapy in the treatment of cancer or auto-immune diseases. To achieve the goal of capturing the diversity, quantifying the abundance of T-cell clones and performing longitudinal comparisons, TCR sequencing has emerged as an approach to monitor T cell responses to therapy and disease progression.

The OTSP pipeline described herein provides a transparent protocol enabling clonality metrics, including evaluating amplification bias specific to each primer pair, and is reproducable across samples. Count variability approximates the negative binomial distribution that can be exploited using estimated NB means. The NB distribution tells us the variation anticipated in ST counts is indicative of a controlled process.

OTSP pipeline results were compared to data generated from the commercial platform by fixing 260 ST-specific scaling factors derived from ST-only measurements (without the presence of genomic DNA). We observed a high concordance between bulk clonality metrics across the two platforms. This observation indicates that the OTSP pipeline can be integrated across batches, samples and platforms, further improving utility of TCR clonality measurements, something not generally possible when using commercial platforms as ST-counts are typically not provided.

OTSP is open and freely available; we anticipate this will allow scaling up the number of measurements substantially. Since we only use computational normalization, rather than addressing differing primer efficiencies at the bench level, the OTSP methodology is also less labor-intensive than previous methods. Notably, OTSP avoids the primer iteration experiments needed to address amplification bias problems.

Since PCR amplification bias is repeatable across samples we have demonstrated the possibility of conducting analyses without the addition of ST beyond the initial calibration. This is achieved by using ST-specific normalization scale factors obtained from independent, ST-only measurements. The idea of using 260 universal ST-specific scaling factors to address amplification bias in mouse models could be explored further, for if deployed this would substantially decrease cost of this methodology as ST are costly. An additional advantage stems from the fact that ST reduce sequencing depth due to competition between genomic DNA and the ST (**Supplemental Figure 6**). We observed that when gDNA amount was kept constant at 600 ng, increasing concentrations of ST led to decreasing detectability of clonotypes, indicating a competition between the ST and clonotypes during the process. This is especially important as indicated by our results revealing that drop-outs can be frequent, even for the most abundant clonotypes (**Supplemental Figure 7**). For example, when the most frequent 0.3% clonotypes from Wild Type spleen tissue were used, we observed that only 119 distinct clonotypes were detected in the 5 replicates, with 32 clonotypes detected in 4 samples, 21 in 3, and 23 in 2, respectively. These drop-outs stem from the stochastic nature of the sampling and could be reduced by increasing the read coverage by not using ST.

## AVAILABILITY

The source code for the analysis is available at https://github.com/burcudem/TCRSeqNormalization

## SUPPLEMENTARY DATA

Supplementary Data are available at NAR online.

## ACKNOWLEDGEMENT

The authors thank the Knight Cancer Institute (NCI P30 CA069533), Bioinformatics, and OHSU Massively Parallel Sequencing shared resources. We are grateful to members of the Coussens Lab, Spellman Lab and Speed Lab for critical discussions, to Meghan Lavoie for technical assistance, and Justin Tibbitts and Teresa Beechwood for research regulatory oversight and animal husbandry.

## CONFLICT OF INTEREST

R.L. Vonderheide is an inventor on a licensed patent relating to cancer cellular immunotherapy and cancer vaccines, and receives royalties for a licensed research-only monoclonal antibody. He is a member of the Lustgarten Therapeutics Advisory working group. R.L. Vonderheide reports having received consulting fees or honoraria from Medimmune and Verastem; and research funding from Fibrogen, Janssen, and Lilly. He is a member of the Lustgarten Therapeutics Advisory working group.

L.M. Coussens is a paid consultant for Cell Signaling Technologies, AbbVie Inc., and Shasqi Inc., received reagent and/or research support from Plexxikon Inc., Pharmacyclics, Inc., Acerta Pharma, LLC, Deciphera Pharmaceuticals, LLC, Genentech, Inc., Roche Glycart AG, Syndax Pharmaceuticals Inc., Innate Pharma, NanoString Technologies, and Cell Signaling Technologies, is a member of the Scientific Advisory Boards of Syndax Pharmaceuticals, Carisma Therapeutics, Zymeworks, Inc, Verseau Therapeutics, Cytomix Therapeutics, Inc., Hibercell., Inc., Alkermes, Inc., and Kineta Inc, and is a member of the Lustgarten Therapeutics Advisory working group.

P.T. Spellman is a paid consultant from Natera and Foundation Medicine Inc.

A.A. Margolin is a Venture Partner at Khosla Ventures and Chief Executive Officer at NextVivo, Inc. and was previously Chief of Data Science at Sema4, Inc.

No potential conflicts of interest were disclosed by the other authors.

## TABLE AND FIGURES LEGENDS

**Supplementary Table 1.**
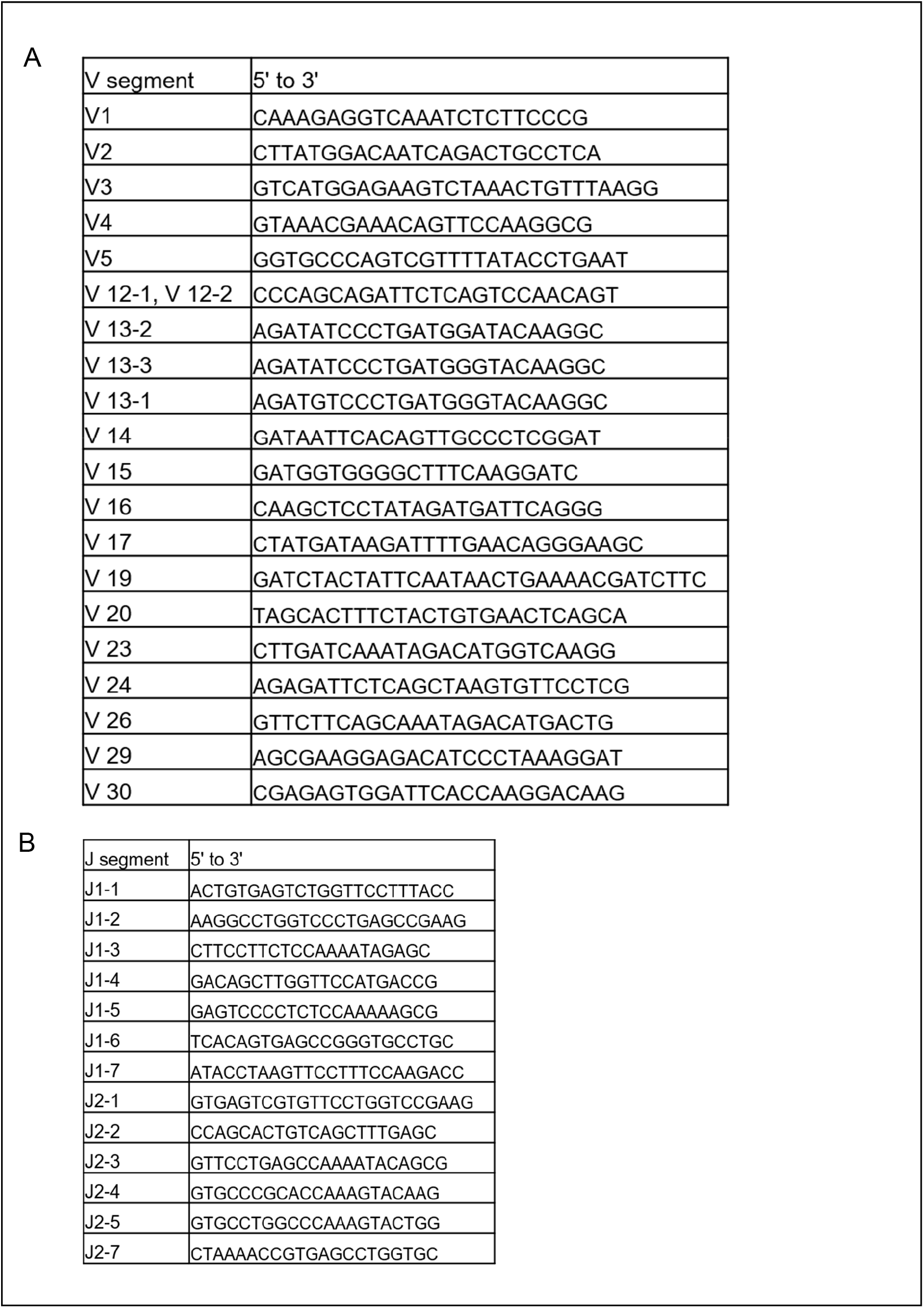
Primer Sequences.

**Supplementary Figure 1.**
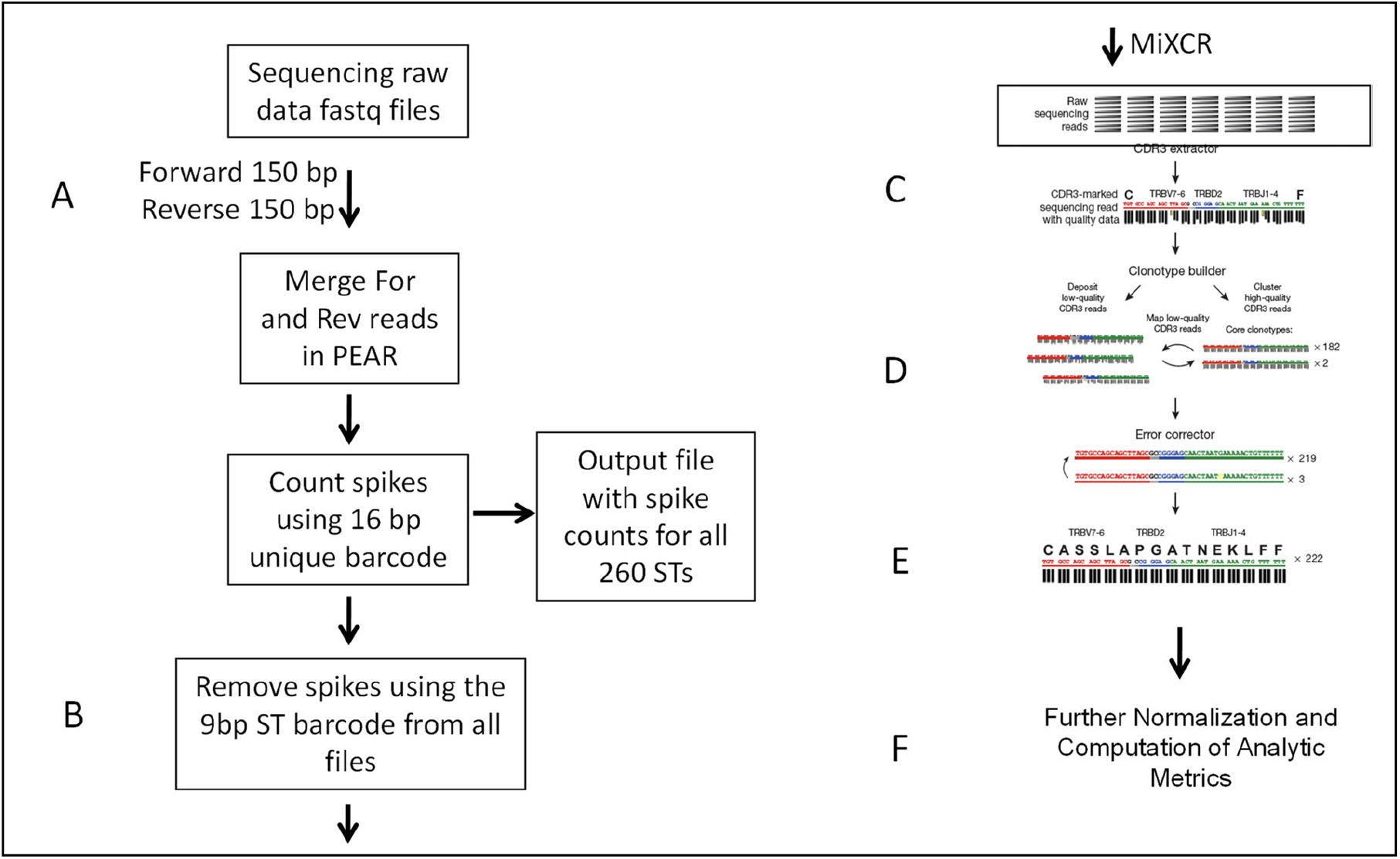
TCR sequencing pipeline schema. Depiction of TCR sequencing pipeline constructed from both extant software tools (configured for use within the pipeline), and dedicated programs written in-house. Multiple samples are processed in parallel, and quality-control checks provide visibility into the pipeline’s operation. Computation is performed on the 5000 cores using ExaCloud computing cluster. Steps in the pipeline include: **A**. Verification of file integrity and merging of paired-end reads; **B**. ST reads are then identified, quantified, and removed; **C**. Clonotypes are then aligned to reference segments, clustered, and quantified; **D**. Clonotype frequencies are then adjusted to account for PCR amplification; **E**. Clonotypes containing frameshifts and stop codons flagged, and output converted for use by visualization and analysis software; **F**. Analytic metrics computed (diversity, clonal expansion and other as applicable) using various tools indicated within the text.

**Supplementary Figure 2.**
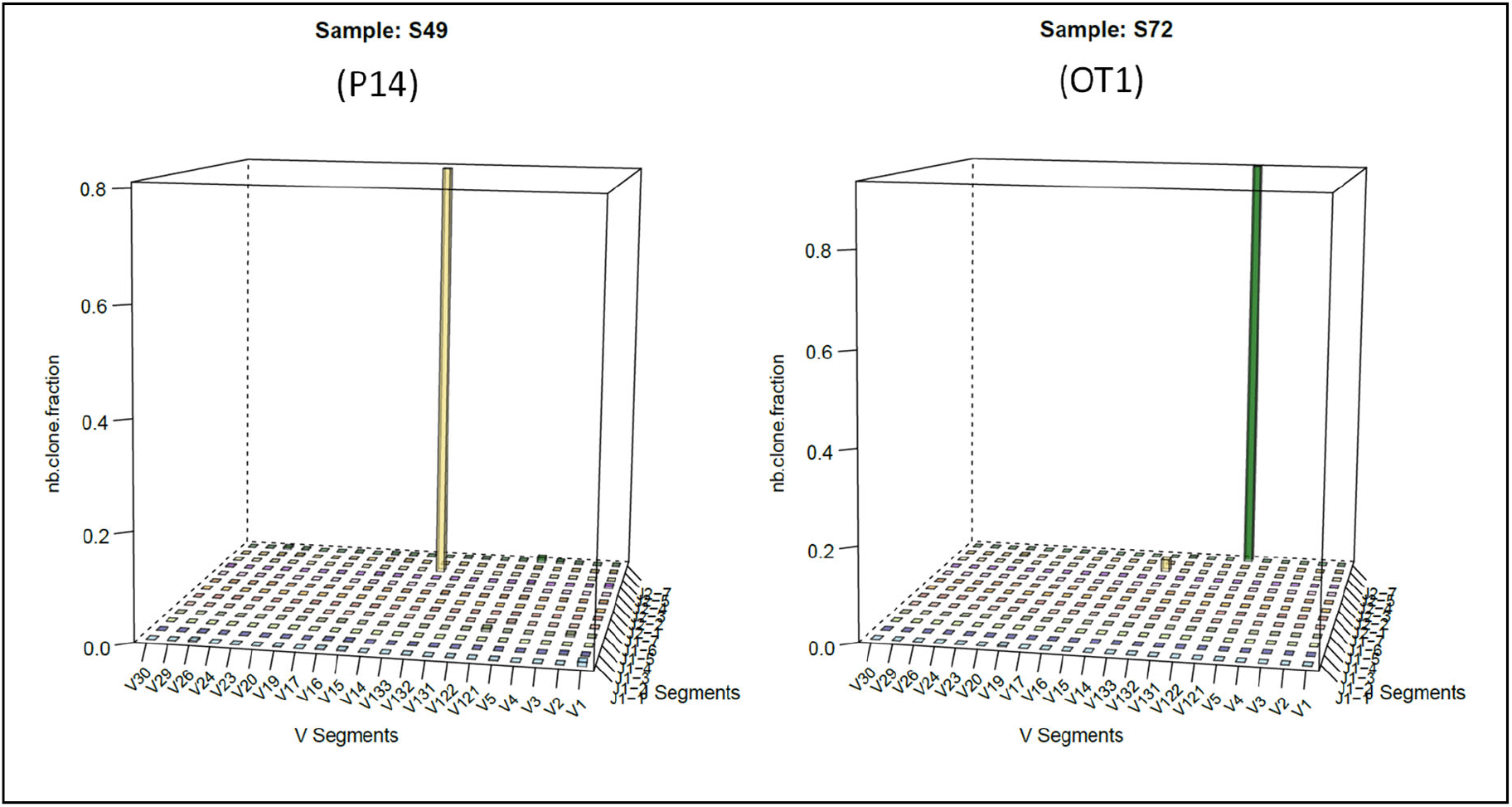
Monoclonal amplification check. Appropriate amplification and identification of clonal TCR segments was verified using OT1 and P14 monoclonal samples, where OT1 was amplified by the primer pair (V12-1,2, J2.7), and P14 amplified by the primer pair (V13-3, J2-4). One example of each monoclonal sample for appropriate amplification is shown.

**Supplementary Figure 3.**
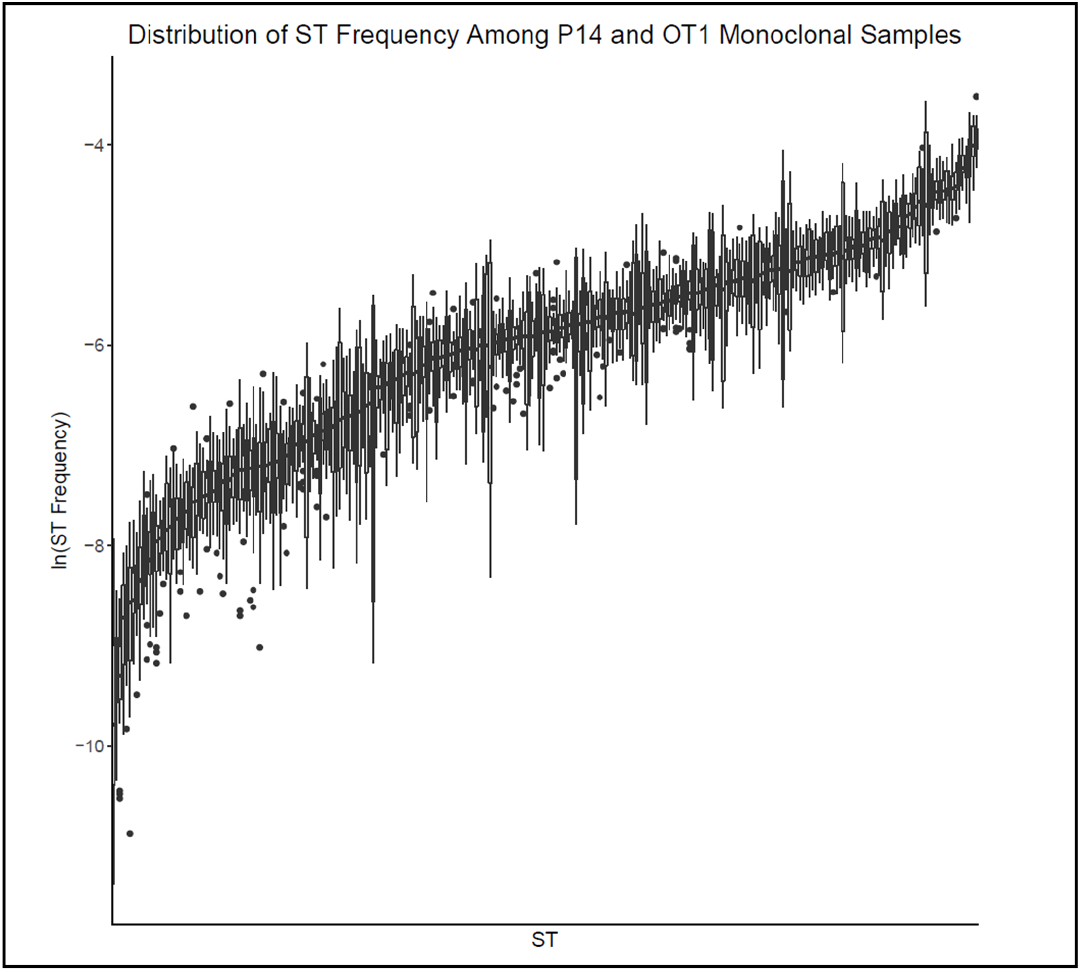
ST Count Distribution in presence of DNA. ST counts were obtained from samples described in the *Transgenic TCR data sets*, where 24 samples of gDNA from P14 and OT1 TCR transgenic mice were processed along with an equimolar mixture of ST. OT1 was amplified by the primer pair (V12-1,2, J2.7), and P14 was amplified by the primer pair (V13-3, J2-4). As with the ST-only samples, when TCR clonotypes were present in the samples along with the ST, the observed variation in the ST counts was caused by the amplification biases of the different primer pairs, rather than by sample to sample variation.

**Supplementary Figure 4.**
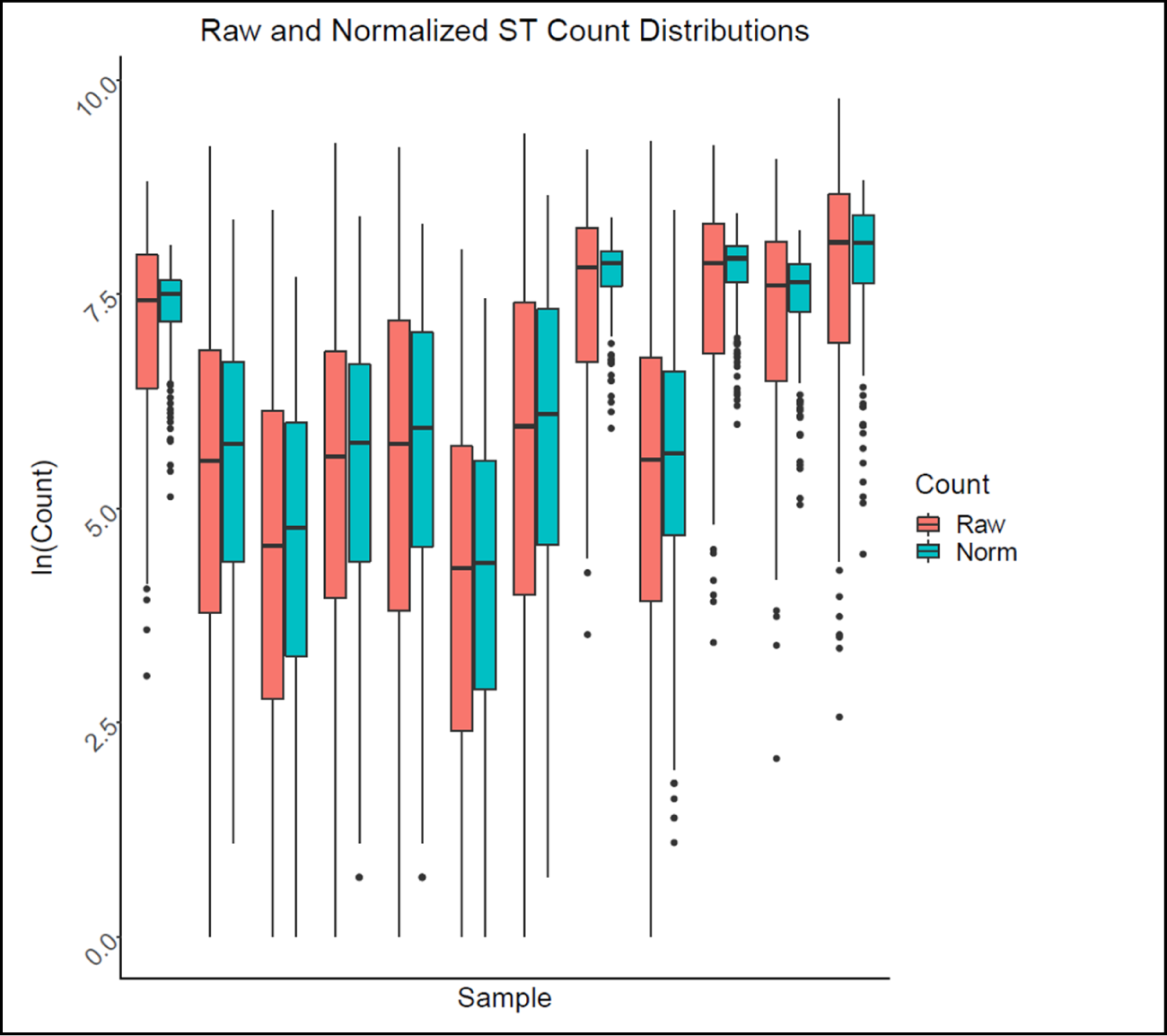
Normalization reduces spread in presence of DNA. gDNA from 12 murine mesothelioma specimens were amplified along with ST (described in the Data Sets section) where the reduction in ST count spread in the presence of DNA before (red) and after (cyan) normalization was plotted.

**Supplementary Figure 5.**
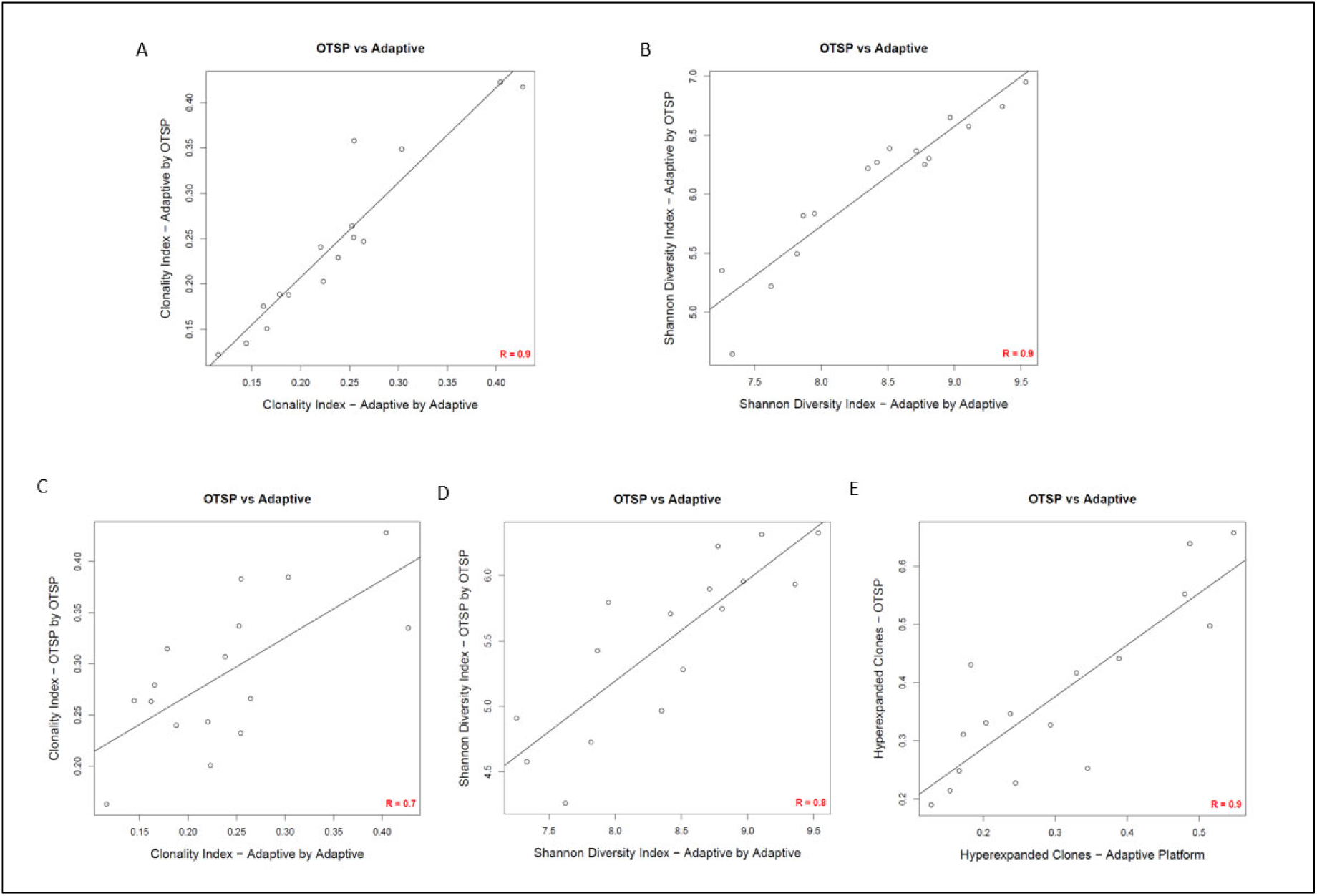
Concordance analysis of T cell repertoire metrics. Concordance analysis comparing commercial and in-house pipelines. gDNA from PDAC tumor samples were evaluated based on a commercial platform (Adaptive Biotech.) where sequences were compared based on output from Adaptive Biotechnology versus OTSP for Clonal index (R=0.9) and Shannon diversity index (R=0.9) **(A-B)**, concordance between the Adaptive Biotech platform versus OTSP for Clonal index (R=0.7) and Shannon diversity index (R=0.8) **(C-D)**, and concordance between the two pipelines for the frequency of hyperexpanded clones **(E)**. p<0.001 for all comparisons with Pearson correlation analysis. Data derived from samples described in the *Byrne et al. data set*.

**Supplementary Figure 6.**
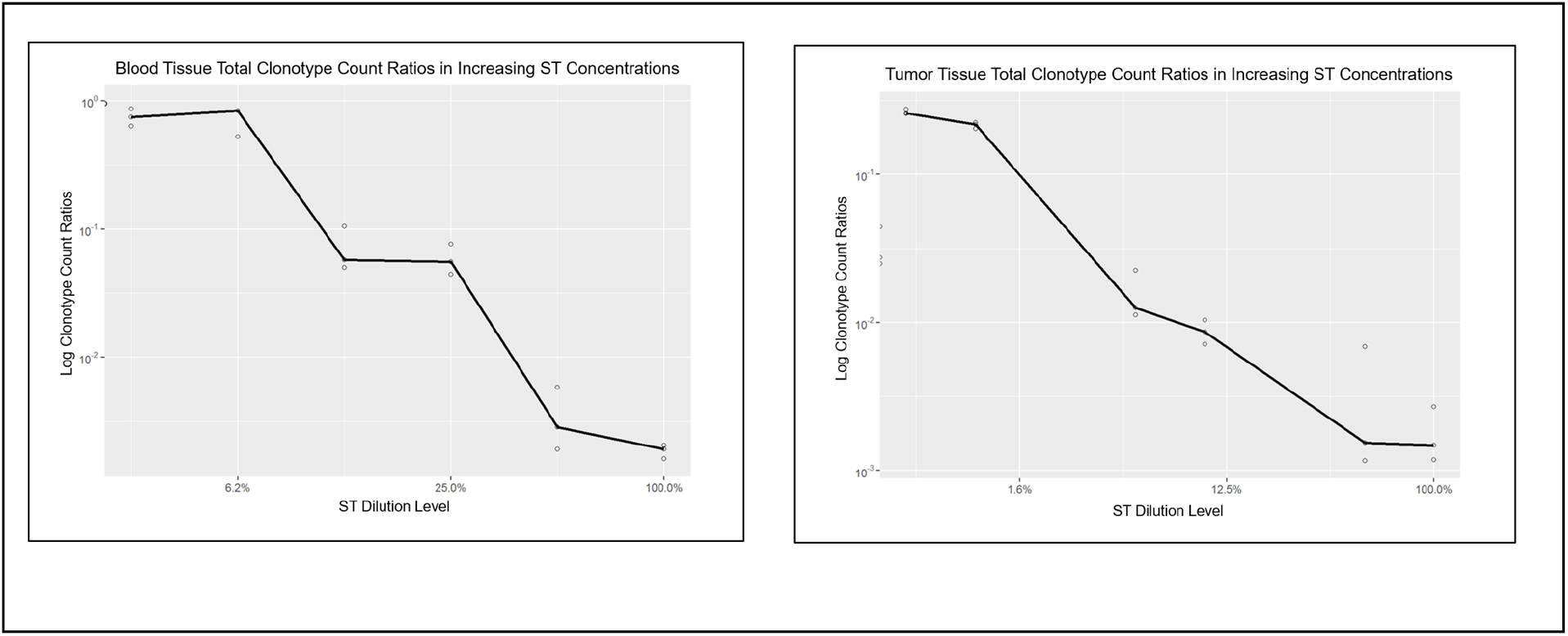
Competition between gDNA and ST during TCR sequencing. Three replicates of 600 ng of gDNA isolated from peripheral blood leukocytes was added to all levels of dilutions (described in the Data Sets section) and for no ST samples for a total of 21 samples plus three replicates of 600 ng of mouse mesothelioma tumor DNA were added to all levels of dilutions and for no ST samples for a total of 21 samples. When the gDNA amount was kept constant at 600 ng, the increasing (relative) concentration of ST lead to decreasing detectability of clonotypes for both type of tissues, showing the competition between DNA and ST occurring during TCR sequencing.

**Supplementary Figure 7.**
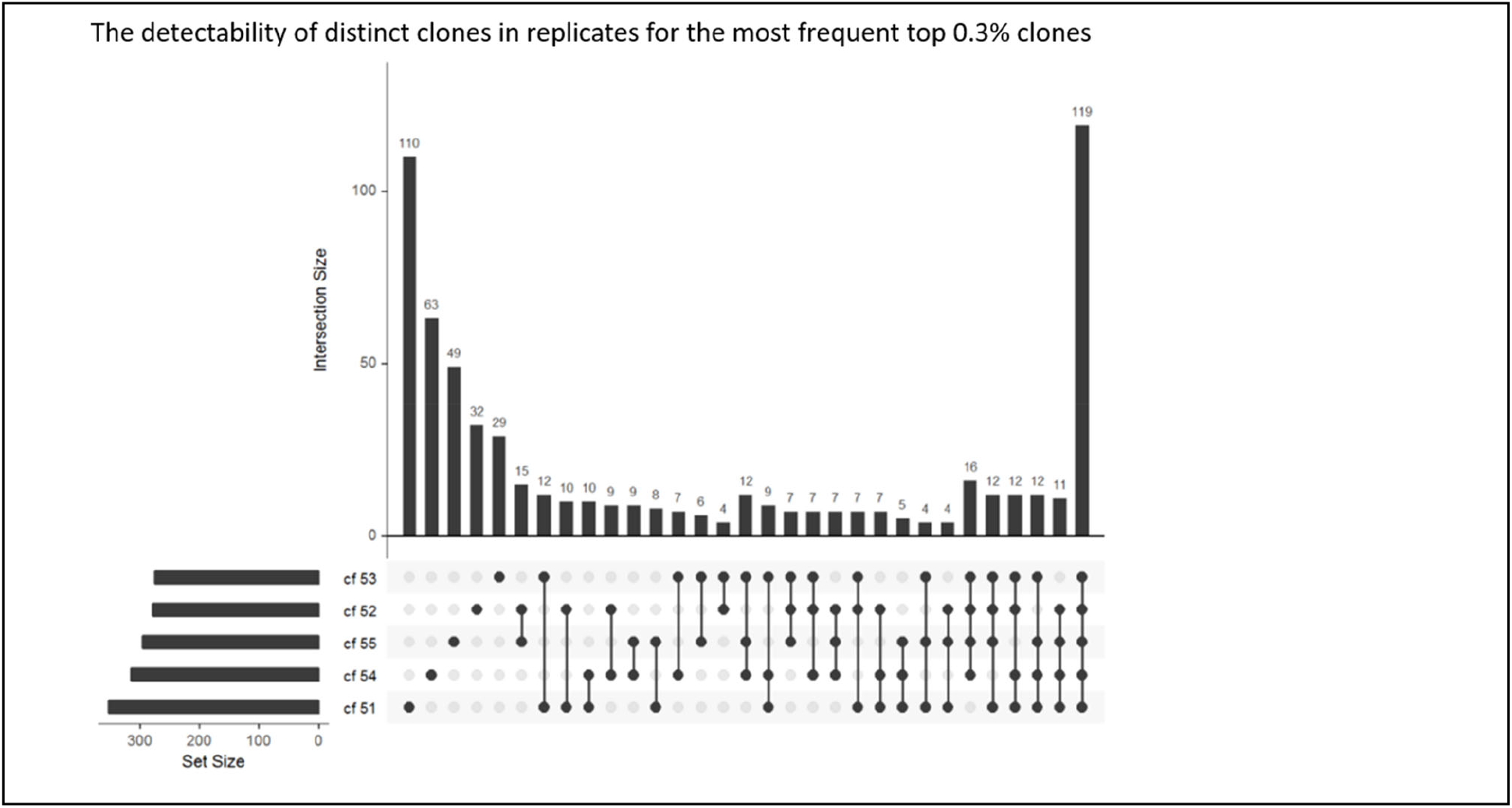
Reproducibility analysis: Drop-outs are frequent even for the top clones. The detectability of distinct clones in five replicates for the most frequent 0.3% clonotypes from wild type spleen tissue is shown. (Data derived from samples described in the *WT spleen data set*.) Vertical bars indicate the frequency of distinct clones detected in replicates, where the number on top of the vertical bars indicates the total number of distinct clones detected in the replicates. On the bottom left, the five replicates (samples cf55-cf55) and the number of distinct clones detected in each replicate is represented by horizontal bars with set size scale. The round dots are black if a particular clone was detected in the corresponding replicate shown at the very left. The connected black dots indicate how many and in which replicates distinct clones were detected out of five replicates. For example, going from right to left, only 119 distinct clonotype were detected in all 5 replicates, 32 clonotypes were detected in 4 samples, 21 in 3 and 23 in 2, respectively. These drop-outs come from the stochastic nature of the sampling.

