## Supplementary Appendix 1 for "An open protocol for modeling T Cell Clonotype repertoires using TCRβ CDR3 sequences"

Suppose that we have three equimolar spike-in only datasets with possibly different concentrations consisting of 20, 10 and 10 replicate libraries, respectively. Denote their ST counts by $\{C_{ij}^{(k)}: i = 1, \dots 260, j = 1, \dots, n^{(k)}\}$, where $i = 1, \dots 260$ labels ST, $j = 1, \dots, n^{(k)}$ labels replicate libraries within datasets, $k = 1, 2, 3$, and $n^{(1)} = 20$, $n^{(2)} = n^{(3)} = 10$.

Our basic assumption is that for any given ST (i.e. primer-pair) the expected values of the counts for are essentially the same, i.e. that we have

$$\boldsymbol{E}\left(C_{ij}^{(k)}\right) = c^{(k)} m_i, i = 1, \dots, 260, j = 1, \dots, n^{(k)}, k = 1,2,3,$$

up to the concentrations $c^{(1)}, c^{(2)}$ , and $c^{(3)}$. Within dataset *k*, natural unbiased estimates of the $c^{(k)} m_i$ are the averages $C_{i\bullet}^{(k)} = \left(n^{(k)}\right)^{-1} C_{i+}^{(k)}$, where $C_{i+}^{(k)} = \sum_{j=1}^{n^{(k)}} C_{ij}^{(k)}$ .

These are maximum likelihood estimates (MLE) under the assumption that all the counts are mutually independent Poisson or Negative Binomial random variables with a common overdispersion parameter for each ST. Our goal here is to show how to combine the three estimates of $m_i$ for any given $i$ taking into account the possibly different concentrations. Without loss of generality we can take $c^{(1)} = 1$, and we will write $c^{(2)} = c$ and $c^{(3)} = d$. Here are two approaches to combining the estimates.

***Assuming independent Poisson or Negative Binomial distributions.*** In this case it is a straightforward calculation to show that the MLE of $\mu_i$ based on all the counts is

$$\widehat{m}_i = (n^*)^{-1} C_{i+}^{(+)} \text{ where } n^* = 20 + 10 \frac{C_{++}^{(2)}}{C_{++}^{(1)}} + 10 \frac{C_{++}^{(3)}}{C_{++}^{(1)}}.$$

This makes sense. We sum *all* the counts observed for ST $i$ and divide that by the sum of the effective number of replicates in each dataset, relative to the concentration for dataset 1.

***Avoiding strong independence and distributional assumptions.*** Here we begin by noting that $log\boldsymbol{E}\left(C_{i\bullet}^{(k)}\right) = log\ c^{(k)} + log\ m_i$, $i = 1, \dots, 260, k = 1,2,3$ and make our goal the linear combination of the three approximately unbiased estimates of $\mu_i = log\ m_i$ , namely the quantities $l_i^{(k)} = log C_{i\bullet}^{(k)}$, $k = 1,2,3$, correcting for the two offsets $\gamma = log\ c$ and $\delta = log\ d$ of the second and third datasets relative to the first, and taking into account the fact that the first dataset has twice their number of observations. A straightforward weighted least squares estimation process leads to the combined estimate of $\mu_i$ as

$$\tilde{\mu}_i = \frac{1}{4}[2 l_i^{(1)} + \left(l_i^{(2)} - \tilde{\gamma}\right) + (l_i^{(3)} - \tilde{\delta})]$$

where $\tilde{\gamma} = \frac{1}{260}(l_{+}^{(2)} - l_{i+}^{(1)})$ and $\tilde{\delta} = \frac{1}{260}(l_i^{(3)} - l_i^{(1)})$. Once we have a combined estimate of $\mu_i = log\ m_i$ , we antilog to obtain our estimate $\widetilde{m}_i$ of $m_i$.

Although the individual ST counts were plausibly negative binomial, they seemed far from independent. As a result, we used the second method to combine the three sets of estimated count means. Recall that in practice, all we need are the estimates of ratios $m_i/m_\bullet$, so that the concentration terms cancel.
