## Supplementary Appendix 2 for "An open protocol for modeling T Cell Clonotype repertoires using TCRβ CDR3 sequences"

Let $C_i$ be the count for spike (primer-pair) *i, i=1,…,n=260.* Our basic assumption is that the $\{C_i\}$ are mutually independent, with $C_i \sim NB(m_i, d_i)$. Thus $\boldsymbol{E}(C_i) = m_i$ and $\boldsymbol{Variance}(C_i) = \boldsymbol{V}(C_i) = m_i + d_i m_i^2$. For the moment, we assume that the parameters $\{m_i, d_i : i = 1, \dots, n\}$ are all known, with $m_\bullet$ and $d_\bullet$ being the average of the $\{m_i\}$ and $\{d_i\}$ respectively.

The normalized counts are $N_i = \left(\frac{m_\bullet}{m_i}\right) C_i$. Clearly $\boldsymbol{E}(N_i) = m_i$ for all *i,* i.e. the normalized values have the same expected value as the original counts.
How variable are they?

Their average is $N_\bullet$ and their empirical variance is $s^2 = (n-1)^{-1} \sum (N_i - N_\bullet)^2$.

We give a lower bound to the expected value of $s^2$, which implies that we cannot use normalization to produce values that are guaranteed to be arbitrarily close together.

***Assertion.*** $\boldsymbol{E}(N_\bullet) = m_\bullet$ , $\boldsymbol{E}(s^2) \geq m_\bullet + d_\bullet m_\bullet^2$ .

***Note.*** The last expression is the variance of an $NB(m_\bullet, d_\bullet)$.

***Proof.*** The equality $\boldsymbol{E}(N_\bullet) = m_\bullet$ follows by averaging both sides of $\boldsymbol{E}(N_i) = m_i$ .
Now $\boldsymbol{V}(N_i) = \left(\frac{m_\bullet}{m_i}\right)^2 \boldsymbol{V}(C_i) = \frac{m_\bullet^2}{m_i} + d_i m_\bullet^2$ , while
$\boldsymbol{V}(N_\bullet) = n^{-2} \sum V(N_i) = n^{-2} \sum \left(\frac{m_\bullet^2}{m_i} + d_i m_\bullet^2\right) = n^{-1} m_\bullet^2 \left\{n^{-1} \sum \left(\frac{1}{m_i}\right) + d_\bullet\right\}$.

The *harmonic mean* of the $\{m_i\}$ is $H = n / \sum \left(\frac{1}{m_i}\right)$, and so $H^{-1} = n^{-1} \sum \left(\frac{1}{m_i}\right)$, and we can write $n\boldsymbol{V}(N_\bullet) = m_\bullet^2 (H^{-1} + d_\bullet)$. .

We now expand $\sum (N_i - N_\bullet)^2$ in a familiar way as

$$\sum (N_i - N_\bullet)^2 = \sum ((N_i - m_\bullet) - (N_\bullet - m_\bullet))^2 = \sum (N_i - m_\bullet)^2 - n(N_\bullet - m_\bullet)^2$$

as the cross term vanishes. Taking ***E*** of both sides, we get

$$\boldsymbol{E} \sum (N_i - N_\bullet)^2 = \sum \boldsymbol{V}(N_i) - n\boldsymbol{V}(N_\bullet) .$$

The rest is algebra. The right-hand side above is

$$\sum \boldsymbol{V}(N_i) - n\boldsymbol{V}(N_\bullet) = \sum \left(\frac{m_\bullet^2}{m_i} + d_i m_\bullet^2\right) - m_\bullet^2 (H^{-1} + d_\bullet) = m_\bullet^2 (n-1)(H^{-1} + d_\bullet).$$

Hence $\boldsymbol{E}\{(n-1)^{-1} \sum (N_i - N_\bullet)^2\} = m_\bullet^2 (H^{-1} + d_\bullet) \geq m_\bullet + d_\bullet m_\bullet^2$ , since $m_\bullet \geq H$,

with equality if and only if the $\{m_i\}$ are all equal.
